# Versican Proteolysis Predicts Immune Effector Infiltration and Post-Transplant Survival in Myeloma

**DOI:** 10.1101/415638

**Authors:** Binod Dhakal, Adam Pagenkopf, Muhammad Umair Mushtaq, Ashley M Cunningham, Evan Flietner, Zachary Morrow, Athanasios Papadas, Chelsea Hope, Catherine Leith, Peiman Hematti, Parameswaran Hari, Natalie S Callander, Fotis Asimakopoulos

## Abstract

High-dose alkylator-based conditioning followed by autologous stem-cell transplantation (ASCT) is a therapeutic mainstay for eligible patients with multiple myeloma. However, post-transplant relapses are common and prognostic biomarkers are scarce. Relapses are characterized by the influx of regulatory myeloid cells and dysfunctional T effectors. We have shown that myeloma-infiltrating myeloid cells produce versican (VCAN), a large matrix proteoglycan with tolerogenic activities. VCAN proteolysis by a-disintegrin-and-metalloproteinase-with-thrombospondin-motifs (ADAMTS) proteases generates versikine, a bioactive fragment (“matrikine”) that regulates Batf3-dendritic cells, known to control CD8+-attracting chemokine networks. Here we demonstrate that intense VCAN proteolysis predicts CD8+ infiltration post-transplant and paradoxically portends significantly inferior survival outcomes. Our data suggest that VCAN proteolysis promotes the influx of CD8+ effectors that are rendered overwhelmingly dysfunctional and/or frankly immunoregulatory (CD8+ Treg) at the tumor site. Thus, complex immunosuppressive circuits orchestrated through VCAN accumulation and turnover generate conditions favorable for myeloma tumor regrowth and point to a readily-assayed biomarker to identify the patients at risk for relapse and early death. The dismal outcomes associated with VCAN proteolysis may be rationally overcome through immunotherapies such as checkpoint inhibition (e.g., anti-TIGIT), tumor vaccines or anti-myeloid (e.g., anti-CSF-1R) approaches.

## INTRODUCTION

The treatment landscape of multiple myeloma (MM), a common cancer of mature plasma cells that secrete immunoglobulin, has been radically transformed with the introduction of proteasome inhibitors and thalidomide analogs (1). Even in the “novel agent” era, autologous stem cell transplantation (ASCT) appears to confer a survival benefit (2). Unfortunately, almost all patients relapse and die of drug-resistant disease (3). Recent advances in immunotherapy have enriched the therapeutic armamentarium with approvals of antibody-based immunotherapies (elotuzumab, daratumumab) and promising experimental results with vaccines and adoptive T cell transfer (4). However, challenges remain: namely, the need to potentiate efficacy, mitigate toxicity and extend benefit for most patients.

The presence of tumor-infiltrating lymphocytes (TILs) is linked to favorable clinical outcomes in many cancers (with notable exceptions characterized by robust Treg infiltration e.g., renal carcinoma (5)), and has predictive value for approaches such as checkpoint blockade (6). In contrast to disseminated hematological malignancies where immunological tolerance is attributable to defective T effector priming and anergy (7), myeloma is characterized by intratumoral effector dysfunction, mirroring solid tumor dynamics (8). Myeloma relapses post-ASCT are heralded by the emergence of dysfunctional (exhausted/senescent) T cells expressing programmed death -1 (PD-1) and T-cell immunoglobulin and ITIM domain (TIGIT) receptors as well as IL-10-secreting myeloid cells (9-13). However, it is not *a priori* clear, which patients will progress after ASCT.

We previously demonstrated that versican (VCAN), a large matrix proteoglycan, is secreted by myeloma-infiltrating myeloid cells and sustains tumor-promoting inflammation and immune suppression (14, 15). Consistent with these findings, VCAN-producing macrophages were shown to expand post-ASCT in myeloma and correlate with minimal residual disease (MRD) persistence as well as clinical relapses (16). By contrast, regulated VCAN proteolysis by ADAMTS proteases correlates with CD8+ infiltration in both solid tumors and newly diagnosed myeloma (17). This proteolysis event generates versikine, a bioactive fragment (“matrikine” (18)), that regulates Batf3-dendritic cell (Batf3-DC) differentiation (17). In addition to their roles in tumor antigen cross-presentation, Batf3-DC are instrumental in effector T cell recruitment through CXCL9/10 chemokine networks (19).

In this article, we show that VCAN proteolysis predicts CD8+ T cell infiltration at day 90-100 post-ASCT when disease response to transplant is routinely assessed and maintenance therapy considered. Intriguingly, intense VCAN proteolysis portends inferior clinical outcomes, including progression-free survival (PFS) and overall survival (OS). These observations support a model in which dysfunctional antigen-specific effectors (9-11), IL-10-induced CD8+ Treg (20), regulatory myeloid cells, matrix proteoglycans and matrikines generate a vicious loop of immunosuppression and sustain conditions that favor relapse. Thus, the unfavorable prognosis conferred by VCAN proteolysis may be rationally reversed through effector potentiation (anti-TIGIT/ anti-PD1 checkpoint inhibition, vaccines) or approaches targeting regulatory myeloid cells (e.g., anti-CSF-1R antibodies).

## RESULTS AND DISCUSSION

Bone marrow biopsies from 35 myeloma patients were collected at Day 90-100 post-ASCT. Table 1 gives the baseline characteristics. The median age was 61 years (range 24-74) and 63% were males. Twenty four percent had high-risk cytogenetics, and 18% had very good partial response or better (≥VGPR) at the immediate pre-transplant evaluation. All patients received high dose melphalan at 200 mg/m^2^ for conditioning. Disease evaluation was done at day 90-100 which showed that 73% of patients were in ≥VGPR. Most patients received maintenance therapy (63%) with no significant differences between VCAN-proteolysis classification groups (Table 1).

**Table 1:**
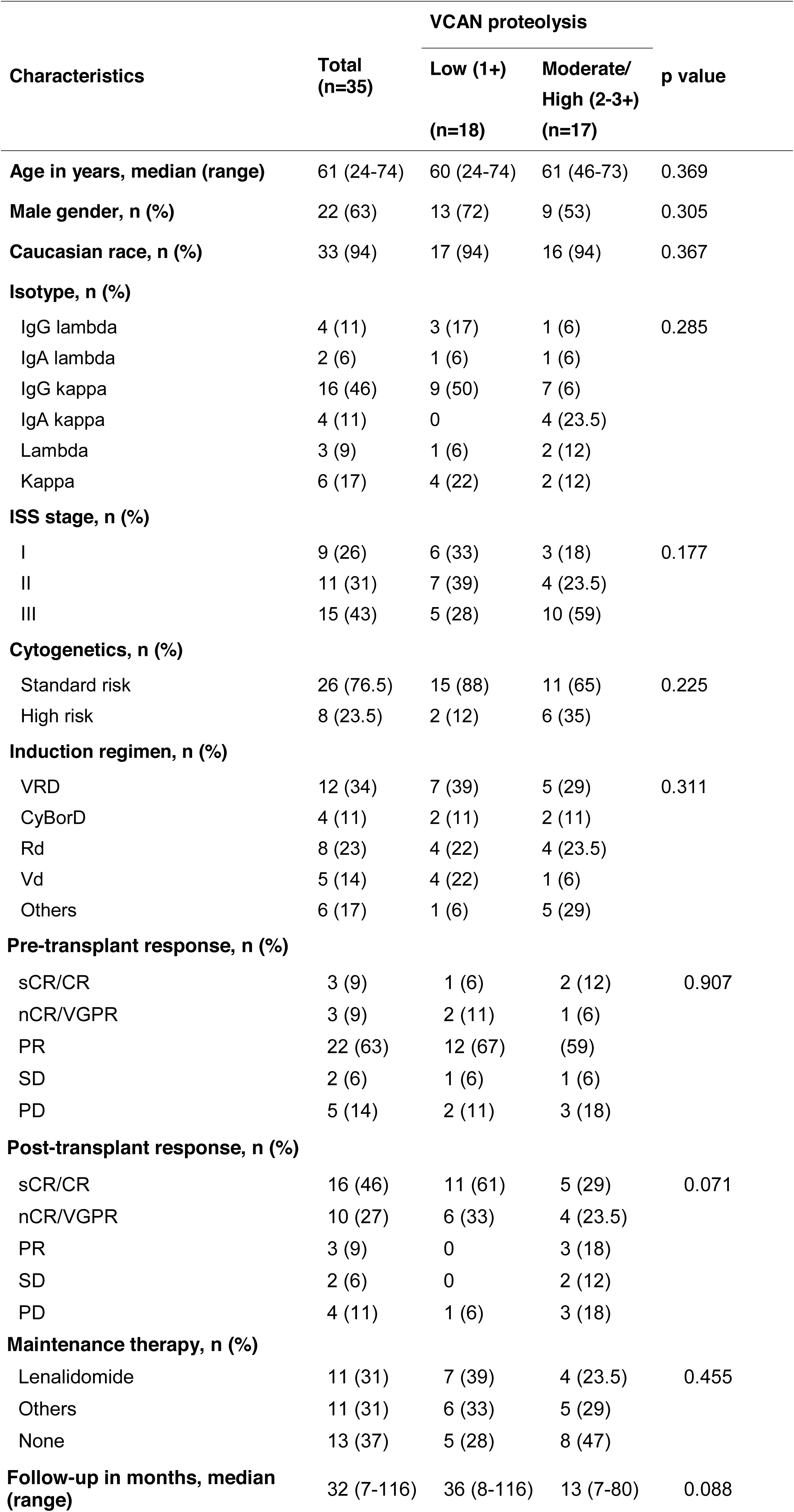
Baseline and post-transplant characteristics of patients

### VCAN proteolysis detection in MM bone marrow biopsies

We stained the core bone marrow biopsies of MM patients against a neoepitope (anti-DPEAAE) generated through VCAN-cleavage at the Glu^441^-Ala^442^ bond of the V1-VCAN isoform (21). DPEAAE constitutes the C-terminal end of the bioactive VCAN fragment, versikine (22). Staining was robust despite prior decalcification of samples, a condition known to adversely impact detection of other immune biomarkers, such as PD-L1, by immunohistochemistry. VCAN proteolysis-low intensity (1+) was present in 18 (51%) patients, while VCAN proteolysis-high intensity (2+ and 3+) was present in 17 (49%) patients (2+ intensity= 11 patients and 3+ intensity= 6 patients). Representative staining patterns are given in Fig. 1.

**Figure 1.**
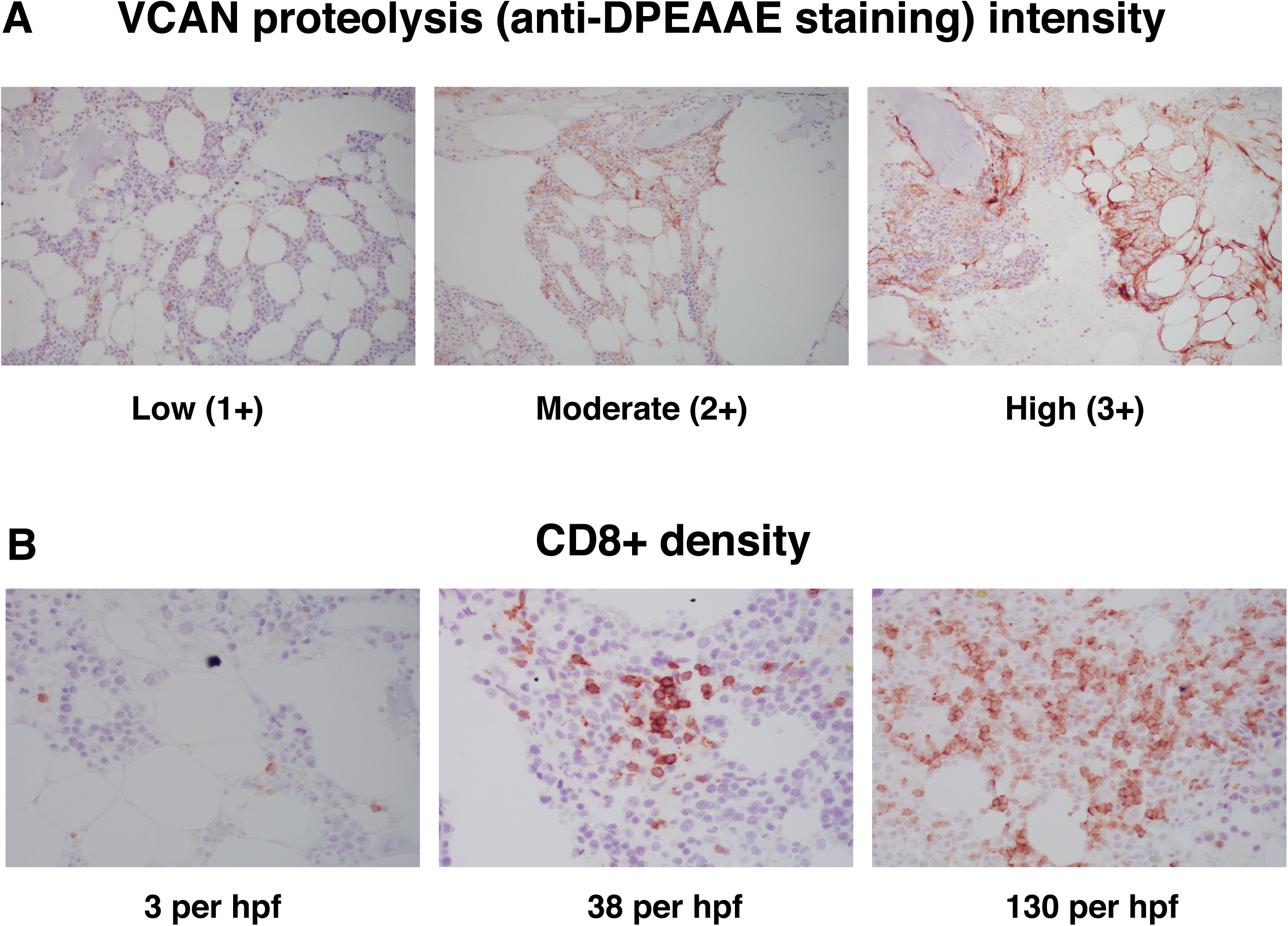
Representative photomicrographs of post-ASCT bone marrow biopsies stained with an antibody detecting VCAN proteolysis (anti-DPEAAE neoepitope) (A) and an anti-CD8 antibody (B). hpf= high-power field.

### VCAN proteolysis correlates with CD8+ T cell infiltration

Core biopsies were stained for the effector T cell marker CD8 and CD8+ counts per high-power field (hpf) were correlated with the VCAN proteolysis status (Fig. 1). There was a statistically significant correlation between VCAN-proteolysis intensity and CD8+ T cell infiltration (VCAN proteolysis intensity 1+: CD8+ count per hpf mean, SD= 17.7 (13.1); VCAN proteolysis intensity 2+: CD8+ count per hpf mean, SD= 27 (14.8); VCAN proteolysis intensity 3+: CD8+ count per hpf mean, SD= 67.3 (43.1); p<0.001) (Fig. 2). On univariate logistic regression, intense VCAN proteolysis (2-3+) was associated with higher CD8+ T cell count (OR, 1.06, 95% CI 1.01-1.12; p=0.027).

**Figure 2.**
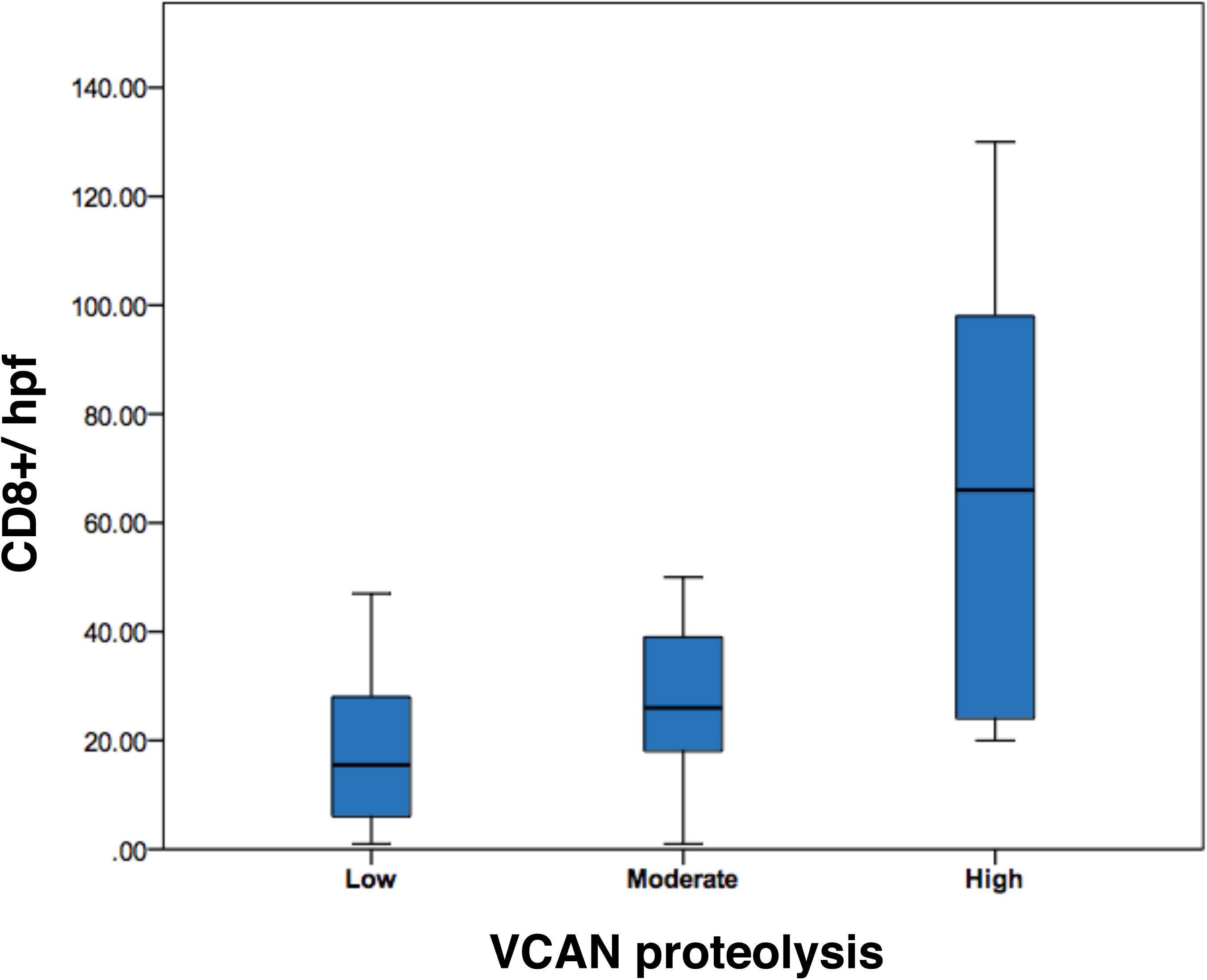
Box-and-whisker plots demonstrating the distribution of CD8+ count per hpf according to VCAN proteolysis intensity.

### VCAN proteolysis status and post-ASCT outcomes

Outcomes were compared between patients with low VCAN proteolysis intensity (1+) and a combined group comprising of patients with moderate and high VCAN proteolysis (2+ and 3+ combined). Overall response rate (ORR) was 83% and 46% patients achieved complete remission. ORR and the depth of response, while superior in patients with low VCAN proteolysis intensity, were not statistically significant (p=0.071, Table 1).

Low VCAN proteolysis compared to moderate/high VCAN proteolysis was associated with better OS (median not reached vs. 13 months, 95% CI 10-16, p=0.003) and a trend for better PFS (median 56 months; 95% CI 26-87 vs. 10 months, 95% CI 1-18; p=0.054) (Fig. 3). In Cox regression analyses adjusted for age and gender, moderate/high VCAN proteolysis compared to low VCAN proteolysis predicted poor OS (HR 4.87, 95% CI 1.33-17.82; p=0.017) and a similar trend was seen for PFS although it was not statistically significant (HR 2.29, 95% CI 0.92-5.68; p=0.074). Higher CD8+ count as a continuous variable was associated with poor OS (HR 1.03, 95% CI 1.01-1.05; P=0.002). In multivariate Cox regression model including VCAN proteolysis status, CD8+ count and cytogenetic risk group, moderate/high VCAN proteolysis compared to low VCAN proteolysis remained independent predictor of poor OS (HR 5.07, 95% CI 1.04-24.76; p=0.045).

**Figure 3.**
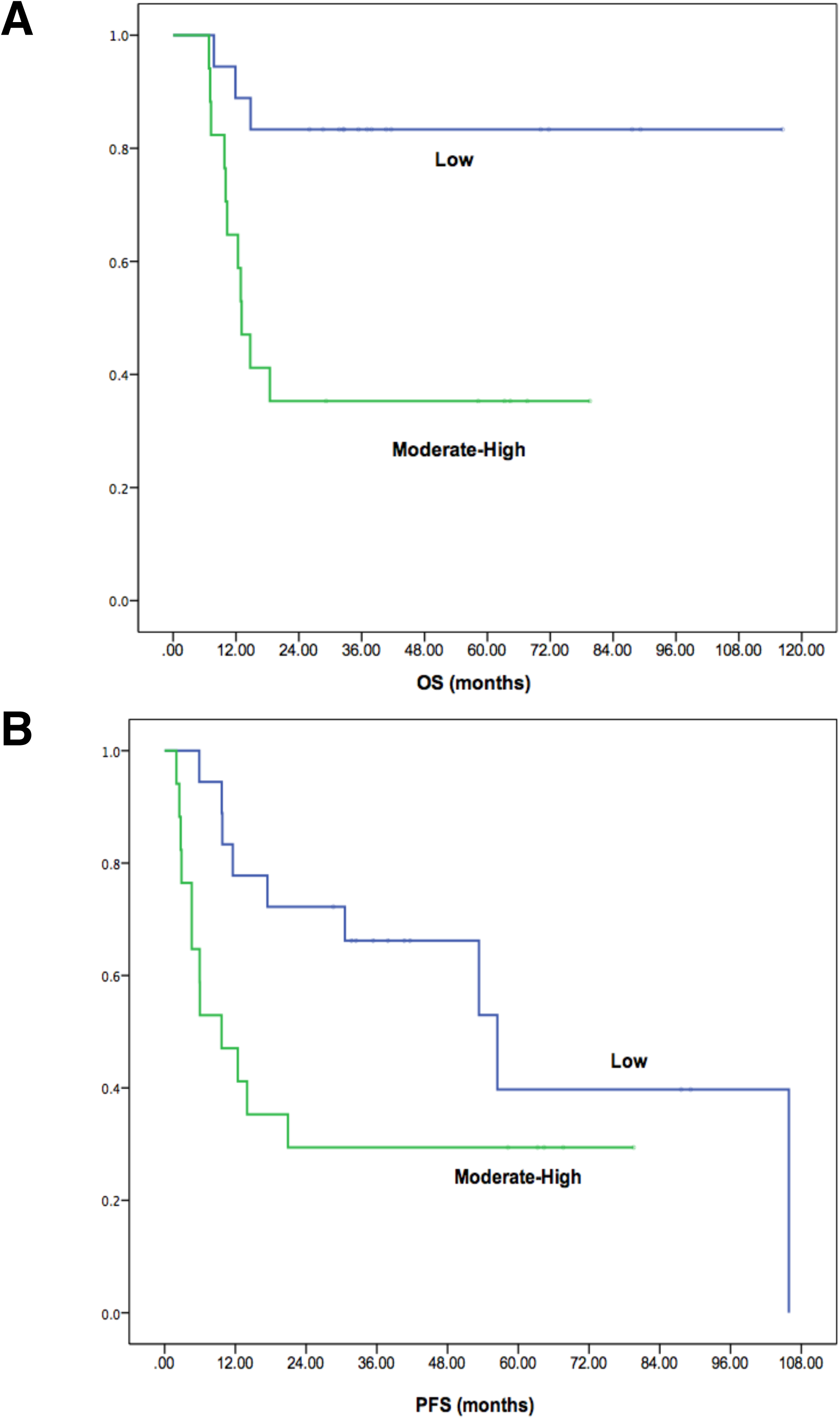
Kaplan-Meier curves demonstrating overall survival (OS, p= 0.003) (A) and progression-free survival (PFS, p=0.054) (B) according to VCAN proteolysis intensity: low (1+) and moderate/high (2+ and 3+ intensity combined).

Patients with low VCAN proteolysis compared to moderate/high VCAN proteolysis had better 2-year PFS (72% vs. 29%, p=0.018) and 2-year OS (83% vs. 35%, p=0.006). In logistic regression analyses adjusted for age and gender, low VCAN proteolysis compared to moderate/high VCAN proteolysis predicted higher 2-year PFS (OR 6.80, 95% CI 1.41-32.76; p=0.017) and 2-year OS (OR 7.64, 95% CI 1.50-39.07; p=0.015). In age-and gender-adjusted multivariate logistic regression model including VCAN proteolysis status, CD8+ count and cytogenetic risk group, low VCAN proteolysis compared to moderate/high VCAN proteolysis independently predicted better 2-year OS (OR 7.77, 95% CI 1.06-56.71; p=0.043) and 2-year PFS (OR 8.48, 95% CI 1.09-66.09; p=0.041).

Multiple myeloma ranks the single most common blood cancer diagnosis with an incidence of 30,770 new cases in the US in 2018 (SEER statistics). Myeloma incidence continues to rise and its association with advancing age and certain environmental toxic exposures is now well established (1). Despite the dramatic improvements in patient outcomes in the last 15 years as the result of “novel agents” (proteasome inhibitors, thalidomide analogs) and advances in stem cell transplantation, myeloma remains incurable for the vast majority of patients.

The typical clinical course is characterized by progressively shorter remissions and relapses with ultimate demise attributable to drug resistance (3). This pattern creates an impetus for the discovery of next-generation novel agents to treat multi-drug-resistant myeloma but also shifts the focus on prolonging early remissions and/or achieving cures through the therapeutic manipulation of anti-myeloma immune responses earlier in the disease natural history. The early post-transplantation period offers an ideal framework for the introduction of immune-activating therapies that aim to generate long-term anti-myeloma memory responses. The constellation of minimal residual disease and immune reconstitution associated with a favorable cytokine milieu underscores the potential of the early post-transplantation period as the optimal timepoint for immunotherapy (11, 23).

Seminal studies dissecting early post-transplant immune effector dynamics have pointed to robust myeloma-antigen specific CD8+ responses that are rapidly rendered dysfunctional in patients who progress: CD8+ dysfunction in myeloma is characterized by CD28 downregulation, CD57 expression, and expression of multiple co-inhibitory receptors, most notably TIGIT and PD-1 (9-12, 24). Moreover, antigen-nonspecific CD8+CD28-CD57+ T-cells were shown to be induced by IL-10 and exert frankly immunosuppressive activity on antigen-specific responses (20). In murine models, IL-10 has been shown to be mostly derived from regulatory myeloid cells, predominantly immature DC and macrophages (10). Targeting myeloid cells e.g., using anti-CSF-1R antibodies or T-cell dysfunction using anti-TIGIT or PD-1 antibodies holds great promise for prolonging post-transplant remissions and generating antigen-specific memory (8-10).

There is clearly a dichotomy in outcomes post-transplant in both human patients and murine systems: some patients will achieve long-term remissions while others rapidly relapse (10). The factors responsible for these disparate outcomes are unknown and prognostic biomarkers are urgently needed. In this work, we tested the hypothesis that VCAN proteolysis may hold value as an indicator of immune surveillance during the early post-ASCT period and correlate with patient outcomes. In earlier work, we showed the accumulation of VCAN, a large matrix proteoglycan, in myeloma tumors (15). VCAN stimulates a TLR2/6-dependent signaling loop that renders dendritic cells (DC) dysfunctional through autocrine IL-6/IL-10 production (25). Intriguingly, the regulated proteolysis of VCAN by ADAMTS proteases is associated with CD8+ infiltration in myeloma bone marrow at diagnosis as well as solid tumor contexts (14, 17). VCAN proteolysis generates bioactive fragments (“VCAN-matrikines”) (18). The VCAN-matrikine, versikine, regulates the differentiation of Batf3-DC (26, 27), from Flt3L-mobilized bone marrow progenitors (17). Batf3-DC, in turn, promote CD8+ effector infiltration, through CXCL9/10 chemokine networks (19).

Our study provides the first set of data ascribing prognostic significance to VCAN proteolysis status. We chose to study post-ASCT bone marrow at the Day 100 timepoint when ASCT-related response is first assessed and maintenance approaches are considered. We observed the paradoxical association of intense VCAN proteolysis, high CD8+ T cell infiltration and dismal post-ASCT survival. This result supports the hypothesis that VCAN proteolytic fragments promote CD8+ infiltration (likely through Batf3-DC chemokine networks) but these CD8+ effectors become dysfunctional and/or frankly immunoregulatory at the tumor site (11). Multiparametric analyses suggested additional negative effects of VCAN proteolysis, *independent of CD8+ T cell status*, alluding to yet unexplored interactions between intrinsically high-risk disease and tumor matrix remodeling. Early (within 2 years of ASCT) progression of myeloma is an extremely powerful prognostic predictive marker of shorter OS (28). Our data indicate that moderate/ high VCAN proteolysis serves as a biomarker at around Day 100 after ASCT that can identify patients at risk of early progression. Early phase clinical data demonstrated the promise of post-ASCT vaccination with a DC-myeloma cell fusion product (29). Despite recent concerns about PD-1 pathway-targeting checkpoint inhibition in myeloma, other checkpoint pathways (e.g., TIGIT) are ripe for clinical testing (9, 10). We propose that risk stratification via VCAN proteolysis staining will pinpoint the patients most likely to benefit from these cutting-edge immunotherapies.

## METHODS

### Patients and Samples

Patient samples were collected with informed consent under a UW-Madison Institutional Review Board-approved protocol (HO07403). Patients underwent ASCT at UW-Madison. Bone marrow core biopsies were collected at day 90-100 to assess disease status post-ASCT.

### Immunohistochemical methods and antibodies

Bone marrow core biopsy samples were fixed, decalcified and paraffin-embedded per standard pathology laboratory procedures. Unstained bone marrow core biopsy slides were deparaffinized and rehydrated using standard methods. Antigen retrieval was carried out in EDTA buffer (CD8 detection) or citrate (DPEAAE). Primary Abs included anti-DPEAAE (PA1-1748A; Thermo Fisher, Waltham, MA) and CD8 (c4-0085-80; eBioscience, San Diego, CA). The anti-DPEAAE neoepitope Ab has been validated previously (21).

### Scoring and analysis of staining patterns

Core biopsy anti-DPEAAE staining intensity was scored for each sample by at least three observers, including a pathologist (AMC), blinded to clinical parameters. Stained slides were examined using Olympus BX43 microscope with an attached Olympus DP73 digital camera (Olympus, Waltham, MA). Immunostaining for anti-DPEAAE was assessed by scoring staining intensity (1 for low/weak, 2 moderate and 3 for strong/intense staining). For CD8+ detection, the number of CD8+ T cells per hpf was calculated using a single area at 400X magnification (10X ocular with a 40X objective). Presence of ≥ 15 CD8+ T cells was considered robust. For clinical outcome analyses, patients were divided into two groups: low VCAN proteolysis (anti-DPEEAAE staining intensity 1+) and moderate/high VCAN proteolysis (anti-DPEEAAE staining intensity 2+3) respectively.

### Statistical analyses

Baseline and post-ASCT characteristics between the low and moderate/high VCAN proteolysis groups were compared using chi-square test, Fisher’s exact test or t-test as appropriate. Correlation of VCAN proteolysis status with CD8+ T cell count per hpf was performed using ANOVA and logistic regression. OS and PFS between the two groups were estimated using Kaplan-Meier method. Cox regression analyses were used to correlate factors with OS and PFS using VCAN proteolysis status as the main effect and hazard ratios (HR) and 95% confidence intervals (CI) were obtained. Logistic regression analyses were used to estimate predictors of 2-year OS and PFS and odds ratios (OR) with 95% CI were ascertained. Data were analyzed using SPSS version 21 (SPSS Inc, Chicago, IL), and p<0.05 was considered statistically significant.

## AUTHOR CONTRIBUTIONS

BD, APagenkopf, MUM, AMC, EF, ZM, APapadas, CH: concept, data collection, experiments, data analysis, editing of manuscript. AMC, CL: pathology expertise and analysis. PHematti, PHari, NSC: clinical care, IRB approval, data analysis, manuscript editing. BD and FA: concept, experimental design, manuscript generation.

## ACKNOWLEDGEMENTS

This work was supported by the Leukemia and Lymphoma Society (6551-18), the American Cancer Society (127508-RSG-15-045-01-LIB), the UWCCC Trillium Fund for Myeloma Research and the NIH (P30CA014520).

## Abbreviations

VRD: (Bortezomib, Lenalidomide, Dexamethasone)
CyBorD: (Cyclophosphamide, Bortezomib, Dexamethasone)
Rd: (Lenalidomide, Dexamethasone)
Vd: (Bortezomib, Dexamethasone)
sCR: (stringent complete remission)
CR: (complete remission)
nCR: (near complete remission)
VGPR: (very good partial response)
PR: (partial remission)
SD: (stable disease)
PD: (progressive disease).

